# A neutrophil-driven inflammatory signature characterizes the blood cell transcriptome fingerprints of Psoriasis and Kawasaki Disease

**DOI:** 10.1101/2020.02.24.962621

**Authors:** Arun Rawat, Darawan Rinchai, Mohammed Toufiq, Alexandra Marr, Tomoshige Kino, Mathieu Garand, Mohammed Yousuf Karim, Seetharama Sastry, Aouatef Chouchane, Damien Chaussabel

## Abstract

**Background:** Transcriptome profiling approaches have been widely used in the investigation of mechanisms underlying psoriasis pathogenesis. In most instances, changes in transcript abundance have been measured in skin biopsies. Fewer studies have examined changes in the blood samples from patients with psoriasis. While changes in the periphery may be less relevant, the blood cell transcriptome analysis presents the distinct advantage of being amenable to comparison across diseases.

**Methods:** Two public psoriasis blood transcriptome datasets were reanalyzed and compared against reference datasets encompassing 16 immune states and pathologies, employing a recently established modular repertoire framework.

**Results:** Although perturbations in psoriasis were relatively subtle in comparison to other auto-immune or auto-inflammatory diseases, consistent changes were observed for a signature associated with neutrophil activation/inflammation. This transcriptional signature most resembled that of subjects with Kawasaki disease.

**Conclusions:** The similarities observed between psoriasis and Kawasaki disease blood transcriptome signatures suggest that immune mechanisms driving the pathogenesis of these diseases may be at least partially shared. This notion is reinforced by case reports describing the development of psoriasis disease in patients with Kawasaki disease. Potential implications for novel therapeutic approaches, including the repurposing of biologic drugs targeting IL17 or its receptor for the treatment of Kawasaki disease are discussed.

## BACKGROUND

Inflammation plays an important role in the host defense mechanism against infections. However, when prolonged or excessive inflammation can causd significant pathology [1–3]. Psoriasis affects ~100 million individuals worldwide [4]. It is a common immune-mediated disease with unique epidermal manifestations caused by the disruption of the skin barrier [5, 6]. Its pathogenesis is often believed to be multifactorial, stemming from elements including environmental triggers, such as sunlight, genetically determined susceptibility, and lately microbiome [1, 6, 7]. Kawasaki disease (KD), also called mucocutaneous lymph node syndrome, is a rare childhood disease affecting mostly children under age five [8, 9]. It presents as an acute self-limited vasculitis that sometimes targets coronary arteries and causes ischemic heart disease [10–12]. The etiology of both psoriasis and KD are unknown. And although their clinical manifestations appear to differ, several case studies reported the co-occurrence of these two diseases in the same subjects, suggesting potentially overlapping pathogenic mechanisms [13–17]. Initial treatment regimens differ for the two diseases, with first line treatment for psoriasis including topically glucocorticoids, vitamin D analogues, phototherapy and other immunoglobulin drugs, like anti-IL17 receptor monoclonal antibody (Brodalumab) [18–20]. While in the case of KD it may consist of initial medication with aspirin and other anti-inflammatory drugs [9, 10, 21]. However, second line therapies in both patients with psoriasis and KD will often consist of the same immunoglobulin-based drugs targeting tumor necrosis factor (TNF)-α (e.g., Etanercept) in case of disease progression.

In order to gain a better understanding of the pathophysiology of psoriasis a number of studies have profiled and compared transcriptome profiles of disease *vs.* healthy skin tissues from the psoriasis patients [22]. In contrast, only a few studies have profiled genome-wide transcript abundance in blood obtained from the patients with psoriasis. The major disadvantage employing blood is that it is in this case less relevant for the study of the pathogenesis. However, it presents the distinct advantage of being amenable to repeat sampling. One blood transcriptional profiling permitting for example the identification of biomarker signatures that may be used for evaluating the therapeutic efficacy of anti-TNF-α treatment [23]. Another advantage of profiling blood of patients with psoriasis is that it permits comparisons with other inflammatory diseases in which the skin is not involved.

Employing a fixed transcriptional module framework, we compared fingerprint signatures derived from publicly available psoriasis blood transcriptome datasets with those derived from a collection of 16 reference datasets spanning a wide range of diseases, including KD [24]. The modular signatures shared between Psoriasis and KD was subsequently subjected to extensive functional interpretation via ontologies, pathways and literature term enrichment, complemented by the use of reference transcriptome datasets. Inferences with regards to disease pathogenesis were drawn and potential implications in terms of strategies for therapeutic intervention in KD discussed.

## METHODS

Re-analyses were carried out using a pre-determined collection of co-expressed gene sets (i.e. “modular repertoire”) that are grouped into modules. The module repertoire panel consists of 382 modules annotated into pathways/ontologies as described by Altman et al (Supplementary Figure 1)[24]. Changes in transcript abundance can be represented at the module-level on a grid on which the position for each module is fixed. Compared to baseline, changes for each module are represented by a blue to white to red gradient, where blue represents under-expression, red represents over-expression and white no changes [24]. The workflow of module repertoire analysis and visualization is implemented via a custom R scripts with the analysis performed with the Rstudio version 1.2 [25] using R version 3.6 [26]. The public datasets of psoriasis blood transcriptome datasets discussed below are available in GEO repository [27] and include:

■ The GSE55201 dataset generated using the Affymetrix U133 Plus (microarray) and consisting of 81 subjects/samples profiles. The study examined the role of IL-17 in ameliorating the systemic inflammation, and the impact on the association of psoriasis complications such as atherosclerosis and consequently ischemic cardiovascular disease [28].
■ The GSE123787 dataset generated using the Illumina HiSeq 2000 (RNA-Seq) platform and consisting of 16 subjects/samples profiles. This study examined the involvement of IL-36 in the extracutaneous manifestations associated with manifestations that are poorly understood in psoriasis [29].
■ The GSE40263 dataset generated using Affymetrix Human Gene 1.0 ST Array and consisting of 30 sample profiles (in six groups). The study examined whether a novel antioxidative peptide can reduce oxidative damage in patients with psoriasis. No publications based on this dataset were found in PubMed.
■ The GSE61281 dataset generated using Agilent 44k Human oligo microarrays and consisting of 52 subject/samples profiles. The study examined the role of innate immunity in the pathogenesis of Psoriatic Arthritis [30].
■ The GSE24759 dataset contributed by Novershtern et al. [31] generated using Affymetrix U133A Genechips and consisting of 211 samples. The samples were collected from 4 to 7 donors in different hematopoietic states. The study focused on diverse yet interconnected regulation of gene expression on the hematopoietic states.
■ The GSE60424 dataset [32] generated using the Illumina RNA-Seq platform consisting of 134 subject/samples profiles. Leukocyte populations were isolated from the blood of healthy individuals and patients with diabetes mellitus type 1, amyotrophic lateral sclerosis, multiple sclerosis (MS pre- and post- interferon treatment) as well as septic patients.
■ A study to understand the molecular effect of etanercept, a TNF-α inhibitor, in PBMC of psoriasis patients [23] were not publicly available (and also no response received after initial communication).
■ The GSE100150 dataset [24] generated using Illumina HT12 v3 Beadarrays made publicly available by our group and consisting of 984 subject/samples profiles (Supplementary Table 1).

## RESULTS

An analysis pipeline relying on a fixed repertoire of blood transcriptome modules developed previously by our group served as a basis for the work presented here [24]. Briefly, module repertoire construction in this context relies on co-expression analyses carried out across 16 reference datasets, which represent 16 distinct immune states (diseases: auto-immune, infectious, inflammatory, as well as physiological states: pregnancy or pharmacological immunosuppression) [24] (Supplementary Table 1). Altogether, these datasets encompassed 985 blood transcriptome profiles and co-expression analyses resulted in a large network, from which 382 densely connected subnetworks were extracted. The genes constituting these subnetworks formed in turn the set of transcriptional “modules” used in our analyses. The R script developed to carry out these re-analyses and generation of plots shown throughout this work is available in a separate publication [24]. Briefly, the analysis workflow consists first, in “annotating” the entire gene catalogue into module-membership (location on grid) information. Second, for each module the number of its constitutive transcripts for which significant differences in levels of transcript abundance were found, is recorded and expressed as a percentage. Finally, results are then mapped as a “module repertoire fingerprint”, where each module occupies a fixed position on a grid (**Supplementary Figure 1**). Changes in transcript abundance measured at the “module-level” in the blood of psoriatic patients in reference to controls is indicated by red or blue spots (showing an increase or decrease in abundance, respectively) (**Figure 1**). These “module repertoire fingerprints” were generated for each of the four datasets identified in GEO. Good concordance was observed in two of the datasets (GSE55201 and GSE123787, shown Figure 2). No clear patterns of changes were observed in the other two datasets (GSE40263 and GSE61281). It may be attributed to the fact that overall the psoriasis signature is subtler than that observed in other inflammatory / infectious or autoimmune diseases. Authors of GSE61281 also confirmed that no differentially expressed genes between cutaneous psoriasis patients and controls were found [30]. Or the fact that a different platform was employed for transcriptome profiling could also be a factor, since our group is well-experienced with analysis of RNA-Seq, Illumina beadarray or Affymetrix Genechip data but less familiar with Agilent arrays used in GSE61281. Or this might also have to do with the small numbers of subjects employed in the dataset (GSE40263).

**Figure 1:**
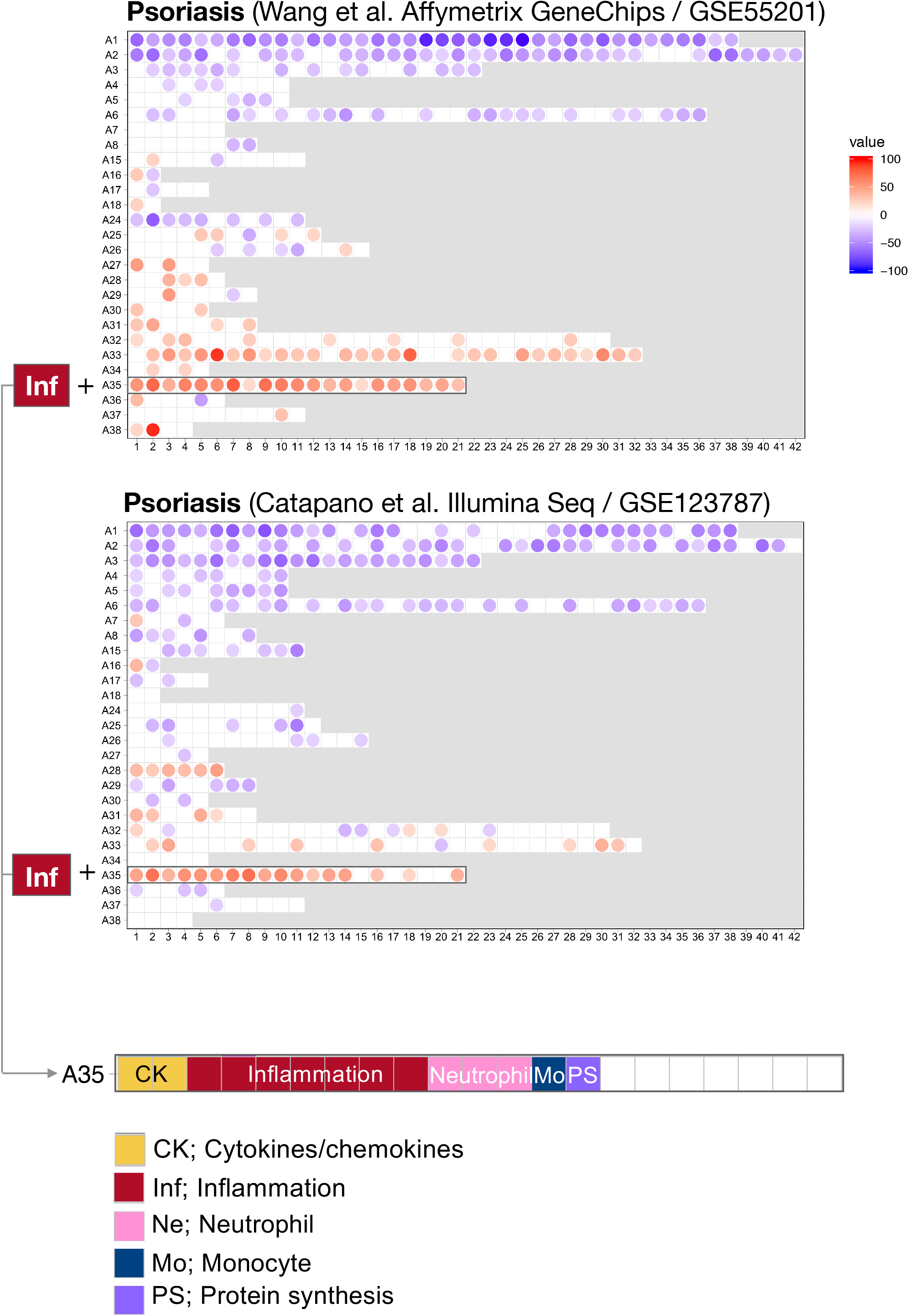
Blood transcriptome fingerprints of psoriasis. Differences in levels of blood transcripts abundance in psoriasis patients and controls are mapped on a grid for the two public datasets, made available by Wang et al. and Catapano et al respectively. Each position on the grid is occupied by a given module (pre-determined set of co-expressed genes). A blue spot indicates a module for which constitutive transcripts are predominantly present at lower levels in patients vs controls. Conversely, a red spot indicates a module for which constitutive transcripts are predominantly present at higher levels in patients vs controls. No spots and white background indicate that there are no changes for the module in question. Grey background color means that there are no changes in the module at this position. Modules are arranged by rows in “module aggregates” and ordered by similarity in expression patterns across a set of 16 disease or physiological states (reference dataset collection). Consistent increase was observed for modules constituting aggregate A35. This aggregate is highlighted on the grid and functional annotations are provided below. Functional annotations for the entire gird is provided in supplementary Figure 1

**Figure 2:**
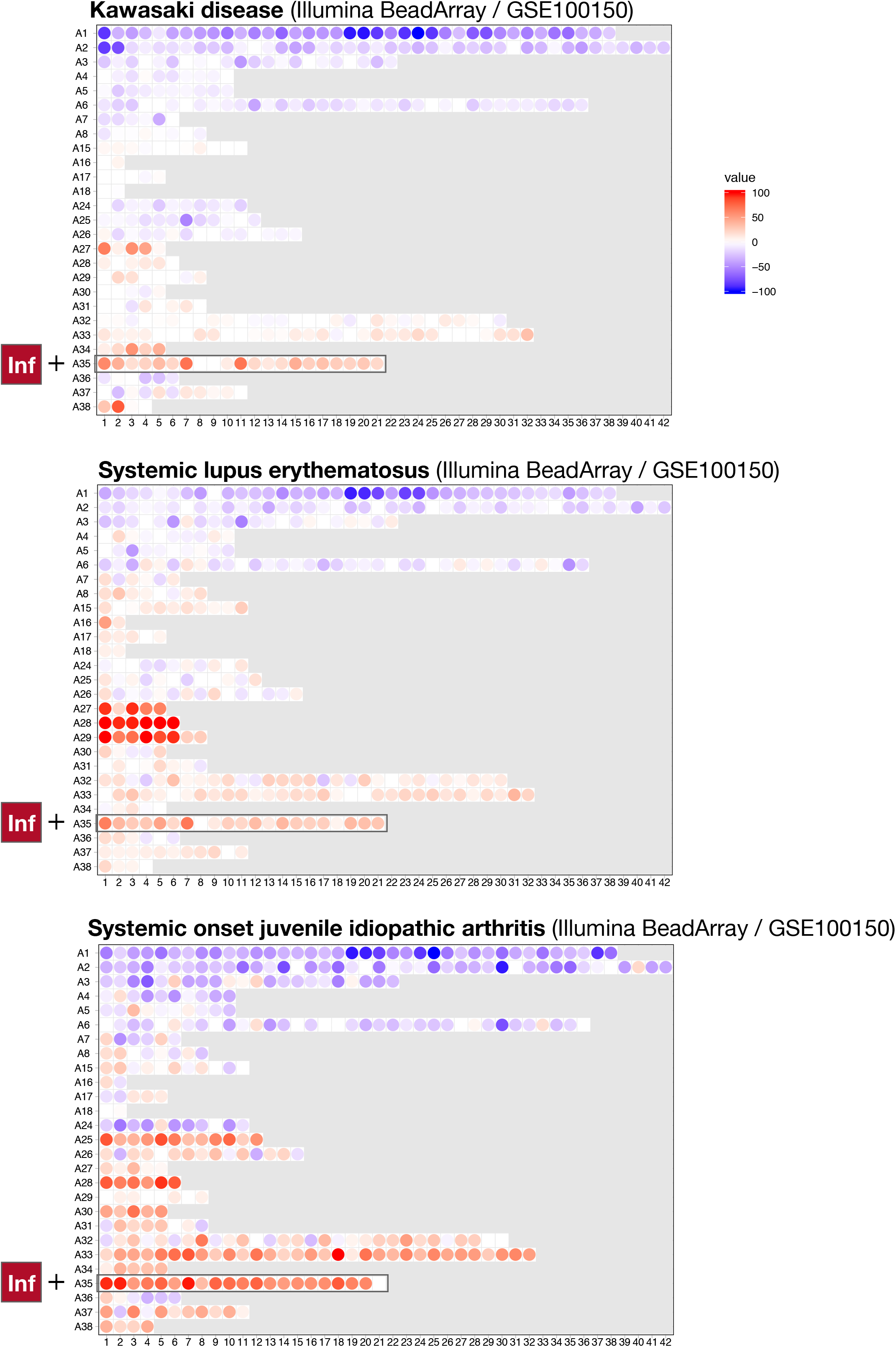
Expression patterns of genes constitutive of A35 modules across whole blood and purified leukocyte populations. Expression levels for genes constituting A35 modules are shown on a heatmap for a reference dataset comprising the profiles of isolated leukocyte populations (GSE60424). Rows represent genes with each cluster of rows representing a module. Columns represent samples. This study compared whole transcriptome signatures of 6 immune cell subsets and whole blood from patients with an array of immune-associated diseases. Fresh blood samples were collected from healthy subjects and subjects diagnosed type 1 diabetes, amyotrophic lateral sclerosis, and sepsis, as well as multiple sclerosis patients before and 24 hours after the first dose of IFN-beta. RNA was extracted from each of these cell subsets, as well as the whole blood samples, and processed for RNA sequencing (Illumina TruSeq; 20M reads).

In the two datasets where signatures could clearly be delineated (GSE55201 and GSE123787), the predominant changes were observed for modules belonging to the following aggregates:

1. A28 in GSE123787 - the modules from this aggregate are functionally linked to the interferon.
2. A35 in both GSE55201 & GSE123787 - the modules from this aggregate are functionally linked to the neutrophil-driven inflammation.

An interferon signature was indeed reported by Catapano et al. and is associated with generalized pustular psoriasis, a more severe form of the disease [1]. It is therefore logical that increases in transcriptional abundance for A28 modules were observed only in this dataset (GSE123787) and not the second one (GSE55201) from Wang et al. [28], since transcriptome profiles of the latter dataset were obtained from patients with the milder and more common form of the psoriasis, compared to the patients associated with the former dataset. The increase in abundance of A35 modules were observed in both datasets, and thus appeared overall as the main component of the blood transcriptome fingerprint associated with psoriasis. Twenty-one module-aggregates comprise the A35 module. Functionally, and at a high level, seven of these modules are associated with inflammation, three with neutrophils, two with cytokine/chemokines one with macrophages and one with protein synthesis (**Figure 1**). The remaining seven modules were not associated with any given functional annotations due to lack of convergence between functional profiling results obtained via different methodologies: reports from gene ontology, pathway and literature keyword enrichment analyses upon which these determinations were made, are available via an interactive presentation: see https://prezi.com/view/7Q20FyW6Hrs5NjMaTUyW/ and **Supplementary Figure 2**. Functional annotations of the module A35 based on the grouping of gene composition of the transcript is provided in **Table 1**. Altogether, dominant functional annotations associated with blood transcriptome of the patients with psoriasis included “neutrophil degranulation”, “inflammation” and “inflammasome”. Some of the genes in these modules which are most recognizable as being involved in inflammatory processes included those coding for inflammasomes components (NLR protein families; found across different modules within this aggregate).

**Table 1:**
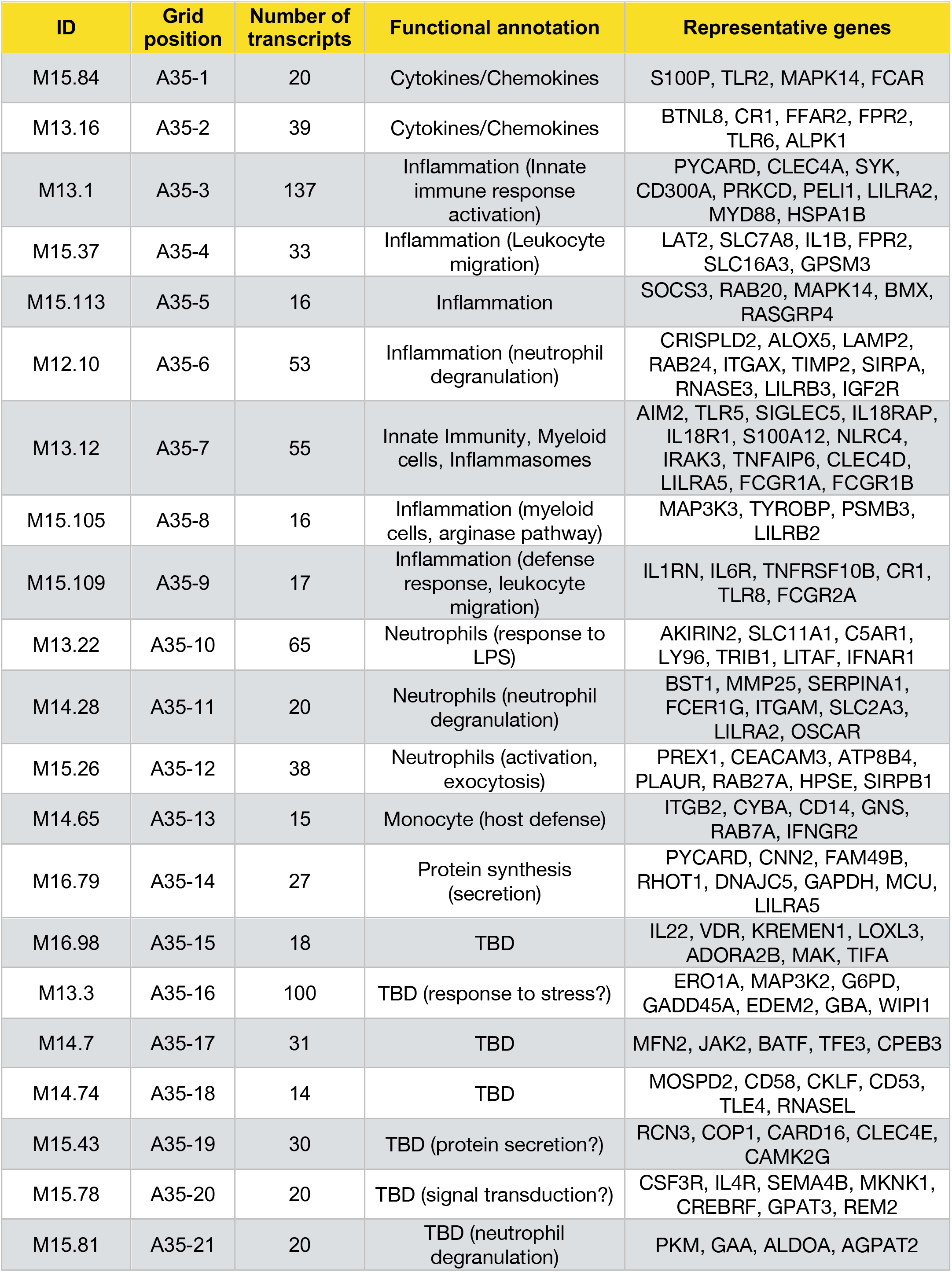
List of modules constituting aggregate A35. Aggregate A35 comprises 21 modules, which are listed in this table along with their respective number of their grid position, number of constitutive transcripts, associated literature keywords and representative genes.

To complement functional analyses, expression patterns of the genes belonging to the module A35 were also obtained for the transcriptome datasets (GSE24759 and GSE60424). Datasets deposited in GEO contributed by Novershtern et al. [31] (GSE24759) and Linsley et al. (GSE60424) [32] were used for this purpose. Both datasets are described in the method section. The first dataset, GSE24759, encompasses a wide range of hematopoietic development states and corresponding heatmaps for each of the 21 A35 modules are accessible via the interactive presentation resource mentioned above (https://prezi.com/view/7Q20FyW6Hrs5NjMaTUyW/). The second dataset (GSE60424) proved particularly informative (Figure 2). It represents RNA-Seq profiles of the leukocyte populations isolated from the blood of healthy individuals and patients with diabetes mellitus type 1, amyotrophic lateral sclerosis, multiple sclerosis (MS pre- and post- interferon treatment) as well as sepsis. Consistent patterns/clusters were observed across the expressed genes constituting A35 aggregate collection of modules, which showed predominant restriction and elevated abundance of neutrophils in leukocytes isolated from septic patients. Altogether, functional profiling and expression profiles observed in reference datasets consistently point towards this signature being associated with neutrophil-driven inflammation.

Next, module repertoire fingerprints obtained for the two psoriasis blood transcriptome datasets (GSE55201 and GSE123787) were compared to a reference collection of fingerprints generated for 16 other diseases [24]. Increase in abundance of A35 modules were observed in blood repertoire fingerprints of systemic lupus erythematous (SLE), systemic onset juvenile idiopathic arthritis (SoJIA) and KD (Figure 3). In the case of SLE and SoJIA, this was one among many other “perturbations” of the blood transcriptome repertoire. This is consistent with the systemic inflammation that characterizes these two diseases. The blood transcriptome repertoire fingerprint of KD was on the other hand more subdued and, like that of psoriasis, dominated by an increase in abundance of A35 module transcripts. This may reflect the fact that the major inflammation is rather more localized in these diseases (skin inflammation for psoriasis and vasculature endothelium (mid-size arteries) for KD). Altogether these results suggest that the extent of perturbations measured at the module level tends to correlate with disease progression from local to systemic inflammation (least perturbed and most perturbed, respectively). As discussed above, the interferon modules A28 along with A35 signature are for instance associated with generalized pustular psoriasis, a severe form of psoriasis. This observation i.e. multiple modules, A28 and A35 in our case, highlights gradient shift from local inflammation towards systemic inflammation also in the context of a given disease.

**Figure 3:**
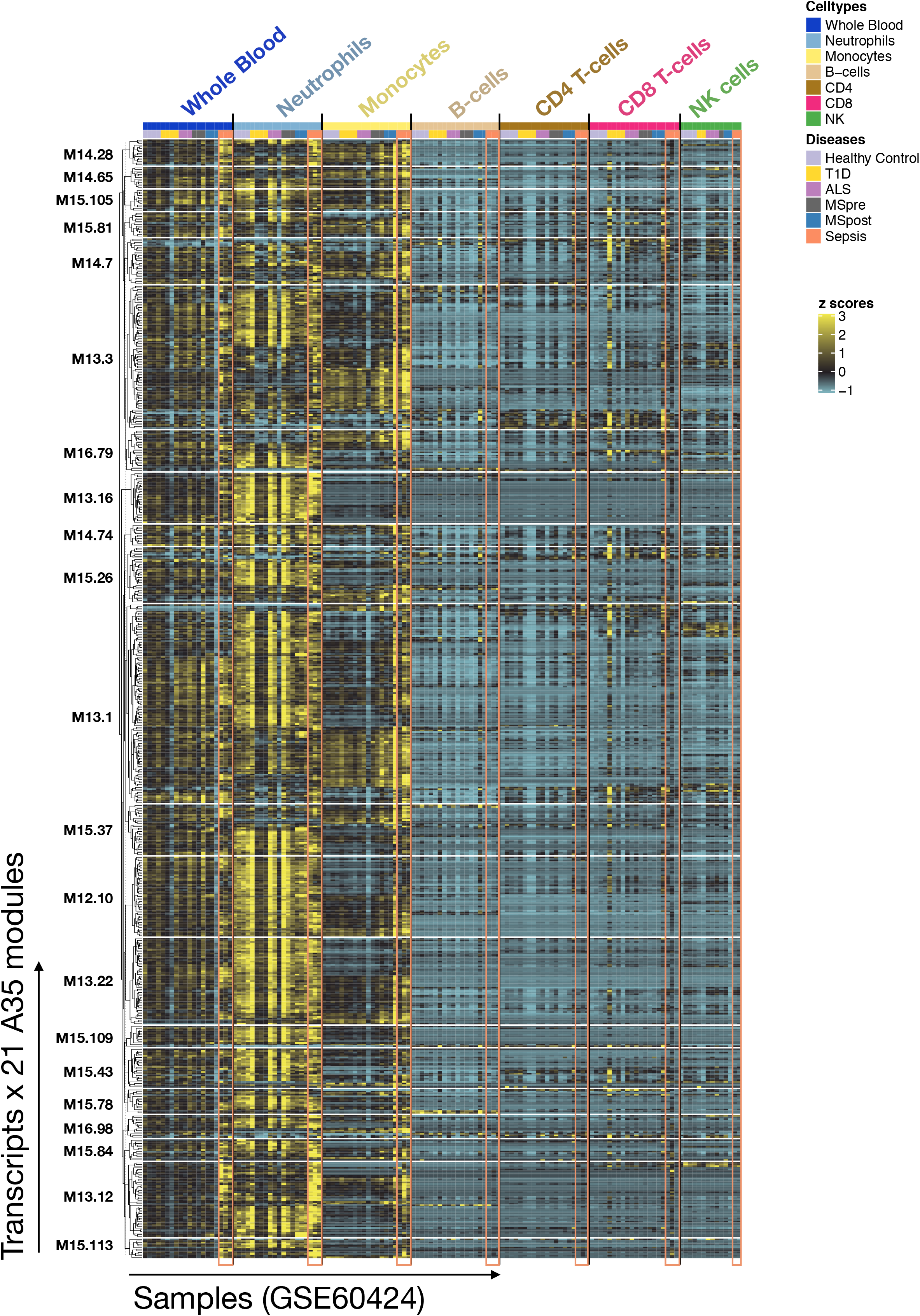
Blood transcriptional fingerprints of other autoimmune or autoinflammatory diseases. Differences in levels of transcript abundance in the blood of patients with Kawasaki disease, systemic lupus erythematous or systemic onset juvenile idiopathic arthritis are mapped on a grid, as described in Figure 1. The modules belonging to aggregate A35 are highlighted on this grid.

## DISCUSSION

First, the comparison of blood transcriptome fingerprints derived from several diseases described above suggests that the blood transcriptome may serve as a means to assess the systemic as well as the localized nature of inflammatory response. This also raises the possibility of using such global profiling approaches to stratify patients based on degree of inflammation, or measure changes over time for a given patient. Actual clinical value, especially against traditional markers of inflammation, such as CRP, would obviously have to be investigated. Secondly, the striking similarities between the two blood repertoire fingerprints identified in psoriasis and KD may suggest that the pathogenesis and/or pathophysiology of these diseases may be driven, at least in part, by similar immune mechanisms. This at least is consistent with a relatively large body of evidence showing that some patients with KD develop psoriasiform eruptions [13–17]. TNF-a, a key inflammatory cytokine in many autoimmune, allergic, inflammatory and infectious diseases, has been shown to play significant role in KD [33] and thus, drugs/medicines targeting this cytokine have been used in its treatment [34]. Of interest here, two reports have shown that psoriasiform eruptions is triggered and/or accelerated in some patients with KD as a result of TNF -α blockade with the approved compound infliximab, [35, 36]. These paradoxical manifestations have also been observed in IBD patients treated with anti TNF-a [37]. These observations, together with the similarities that have between noted earlier between blood transcriptome repertoire fingerprints, suggest a possible overlap of the mechanisms underlying pathogenesis of these two diseases. We inferred that IL17-mediated inflammation could be such common pathogenic pathway. Indeed, for one its role is well-recognized in psoriasis and drugs blocking its activity have been found to be effective and have been approved for therapeutic use. Secondly, this cytokine may also account for the neutrophil-driven inflammatory signature reported here as being elevated in Psoriasis and KD patients since it is a known driver of neutrophil development, maturation and activation. Finally, independent reports having linked Th17 responses to KD also contribute to support this notion [38–41]. A corollary for this observation is that biologic drugs targeting IL17 may also prove efficacious for the treatment of KD, which is something that to our knowledge may not have been tested [based on a search performed on Jan’2020 in Pubmed and clinicaltrials.gov].

## CONCLUSIONS

Despite inherent limitations, especially in diseases where pathology predominantly occurs at local tissue sites, blood transcriptome profiling approaches may be of some use for investigations of diseases such as psoriasis. For one, samples can be obtained for transcriptome profiling with minimal risk and the approach is amenable to serial collection at multiple time points for monitoring applications. And as we have shown here another advantage stems for the availability of “contextual data” for comparative blood transcriptome fingerprint analysis. It is on this basis that we were able to contrast systemic responses across a range of disease states. And it is also on the basis of parallels that could be drawn between the fingerprints of KD and psoriasis that we are able to infer a potential benefit of repurposing anti-IL17 therapy for the treatment of KD.

## ACKNOWLEDGEMENTS

We are thankful to fellow researchers who made public the transcriptome datasets employed in this study.

---

**Supplementary Figure 1:**
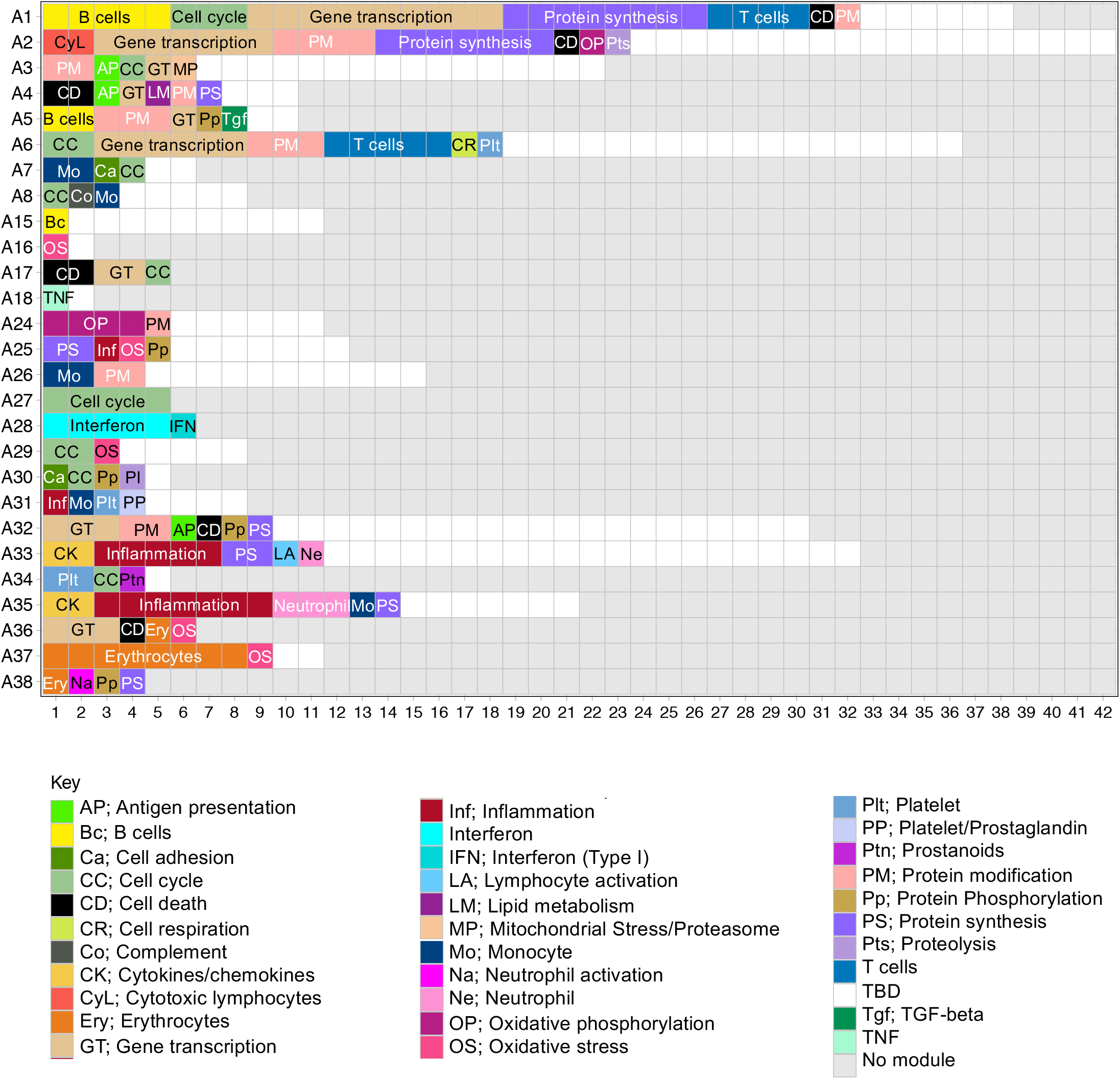
Module annotation grid. This figure provides complete annotation for the grid employed to map differences in transcript abundance between cases and controls shown in Figure 1. Each position on the grid corresponds to a different module. Each of 382- module is constituted by a set of transcripts found to be co-expressed across a range of disease and physiological states. In turn, modules are arranged in rows based on expression similarities. Gene ontology, pathway or literature keyword enrichment analyses provides the basis for attribution of biological functions to the modules. This is indicated by a color code and abbreviations below the grid. An interactive presentation is available which permits the exploration of functional enrichment results and expression patterns for A35 modules is as follow: https://prezi.com/view/7Q20FyW6Hrs5NjMaTUyW/

**Supplementary Figure 2:**
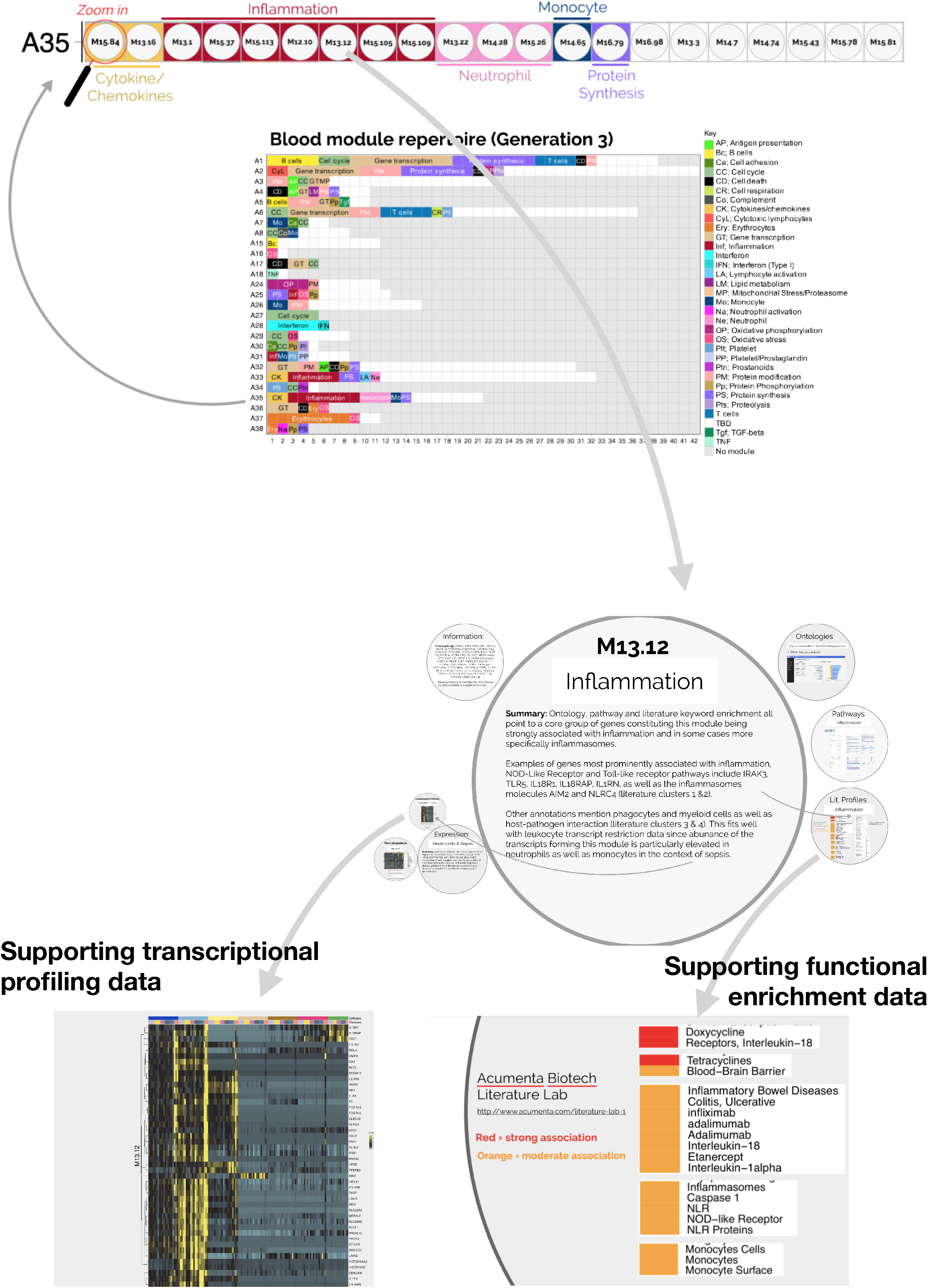
Interactive presentation providing access to supporting transcriptional profiling and functional enrichment data for modules constituting aggregate A35. An interactive presentation is made available to readers and allows exploration of the modules constituting the aggregate A35. A gene list is provided for each module, along with ontology, pathway or literature term enrichment results as well as transcriptional profiling data for reference transcriptome datasets (circulating leukocyte populations, hematopoiesis). It also provides a summary of those findings. The interactive presentation is available via: https://prezi.com/view/7Q20FyW6Hrs5NjMaTUyW/. It provides zoom in/out functionalities for close up examination of the text and figures which are embedded in the presentation.

